# Ancient tandem duplication of *CYCLOIDEA* is associated with the evolution of capitulum development in Asteraceae

**DOI:** 10.64898/2026.08.30.748160

**Authors:** Deng Zhang, Yuan Gao, Yuetong Qian, Yanyan Sun, Jing Zhang, Yuan Liao, Yuhua He, Sumei Chen, Fadi Chen, Baoqing Ding

## Abstract

Asteraceae is one of the largest and most diverse families of flowering plants. Much of their evolutionary success is attributed to their unique inflorescence structure—the capitulum. Despite their profound adaptive and taxonomic significance, the evolutionary genetic mechanisms underlying Asteraceae capitulum diversification remains poorly understood. We integrated comparative genomic analyses of 35 Asteraceae genomes with developmental transcriptome profiling of ray and disc florets. Our results uncovered a deeply conserved *CYC2e*–*CYC2c*–*CYC2d*–*CYC2g*–*CYC2b* tandem gene cluster that is prevalent across the Asteraceae family. We demonstrated that this tandem cluster arose during the early evolutionary diversification of Asteraceae. Subsequent lineage-specific gene retention and loss events have reshaped the composition of this cluster across different Asteraceae lineages. Notably, key members of this cluster, particularly *CYC2c*, *CYC2d*, and *CYC2g*, are preferentially retained in lineages with ray florets, which is consistent with their roles in regulating the ray floret development. Collectively, our findings establish a mechanistic link between the evolutionary divergence, spatiotemporal expression differentiation of the *CYC2* tandem cluster as a key genetic driver underlying the morphological diversification and evolutionary success of Asteraceae.

## Dear Editor

Whole genome duplication is a recurrent theme during plant evolution (Chen et al., 2026). It is considered to have greatly contributed to the evolution of plant morphological and physiological diversity (Soltis and Soltis, 2016). Similar to genome scale duplication, segmented or tandem duplication has constantly provided genetic material for evolutionary innovations and plant adaptation to adverse environments (Hanada et al., 2008; Panchy et al., 2016; Yang et al., 2025) and have facilitated the development of key agronomic traits in domesticated crops (Alonge et al., 2020).

Asteraceae is one of the largest and most diverse families of angiosperms. This extraordinary diversity encompasses a wide range of life forms, from annual and perennial herbs to shrubs and trees, and is associated with exceptional ecological breadth across nearly all terrestrial habitats (Palazzesi et al., 2022). Their evolutionary success has largely been attributed to their unique inflorescences, capitula that mimic solitary flowers but are typically aggregates of multiple florets of different forms (Zhang and Elomaa, 2024). The capitulum typically consists of two distinct floret types: bilaterally symmetrical ray florets and radially symmetrical disc florets, which together contribute to reproductive specialization and pollinator attraction (Andersson, 2008; Stuessy et al., 1986). Beyond their ecological success, the family holds considerable ornamental importance. It includes numerous iconic horticultural species, such as *Chrysanthemum*, *Gerbera* and *Bellis*, which are widely cultivated for their diverse inflorescence architectures, vibrant pigmentation (Song et al., 2023; Su et al., 2019).

Phylogenetic studies suggest that the ancestral Asteraceae most likely possessed a solitary, homogamous, discoid capitulum with isomorphic bisexual florets (Panero and Funk, 2008; Pozner et al., 2012; Zhang et al., 2021a). The majority of extant diversity within Asteraceae is inferred to have arisen from a major radiation during the early Cenozoic, followed by repeated diversification events, particularly during the Paleogene, which further promoted its global expansion and ecological success (Barreda et al., 2015; Mandel et al., 2019). Concomitant with the radiation of the Asteraceae, the family exhibits extensive variation in ploidy, chromosome number, and genome size, reflecting complex evolutionary processes such as polyploidization and hybridization (Huang et al., 2016; Moore-Pollard et al., 2025; Vitales et al., 2019; Zhang et al., 2021a; Zhang et al., 2024). Developmental genetic analysis has uncovered numerous regulators were involved in the capitulum development (Bello et al., 2017; Broholm et al., 2008; Chapman et al., 2012; Chen et al., 2018; Elomaa et al., 2018; Gurung et al., 2024; Shen et al., 2021; Tahtiharju et al., 2012; Uimari et al., 2004; Zhang et al., 2021b; Zhao et al., 2016; Zoulias et al., 2019). Despite their ecological, evolutionary, and economical significance, the molecular origin of Asteraceae capitulum development remains largely obscure.

To explore the genomic mechanism underlying the capitulum diversification, we performed comparative genomic analyses using 35 chromosome-level Asteraceae genomes with diverse capitulum morphologies (Figure S1A and Table S1), representing four major lineages (Panero and Funk, 2008), including Mutisioideae (n = 1), Carduoideae (n = 5), Cichorioideae (n = 4), and Asteroideae (n = 25). Given that the discoid capitulum is considered as an ancestral state, we hypothesized that ray floret evolution involved the acquisition of novel genetic components or regulatory modifications from ancestral disc floret developmental programs. Specifically, we reasoned that these novel genetic components may present in the Asteroideae ancestor with ray and disk florets, but absent from the Carduoideae ancestor with disk florets only.

We first inferred orthologous relationships among the 35 Asteraceae species using FastOMA (Majidian et al., 2025) and reconstructed the ancestral gene complements of the most recent common ancestors of Carduoideae and Asteroideae in pyHam (Train et al., 2019). By comparing these ancestral genomes, we identified 6,119 hierarchical orthologous groups (HOGs) that were present in the Asteroideae ancestor but absent from the Carduoideae ancestor. Gene Ontology (GO) enrichment analysis showed that these Asteroideae-specific HOGs were significantly enriched in developmental and regulatory processes, including meristem growth, floral organ identity, cell differentiation, and hormone signaling (Table S2). Such lineage-specific developmental regulation may underlie the remarkable diversity of capitulum and floret morphologies in Asteroideae. To further refine candidate genes associated with ray floret evolution, we integrated transcriptomic datasets from Asteroideae species with heterogamous capitula, with the expectation that these genes should be differentially expressed between the ray and disk florets. We searched public transcriptome resources, and identified two species with both ray- and disc-floret transcriptomes available: *Helianthus annuus* and *Chrysanthemum indicum*. We identified genes upregulated in ray florets in both species and intersected them with the Asteroideae specific HOGs reconstructed from ancestral genomes. This analysis yielded 14 candidate HOGs potentially associated with ray floret evolution (Figure S1B). GO enrichment analysis of the 14 candidate HOGs revealed significant enrichment in developmental processes, including regulation of meristem growth, plant organ formation, branching morphogenesis, and cell cycle progression (Figure S1C). Pfam annotation showed that two HOGs encode proteins containing a conserved TCP domain (Table S3). It is interesting to note that two auxin inducible genes were also identified, which have recently been shown to be associated with the ray and disk floret development in the Chrysanthemum cultivars (Jia et al., 2026). Phylogenetic analysis further revealed that these genes clustered within the *CYC2g* and *CYC2d* lineages, respectively (Figure S2). These results suggest that members of these *CYC2* subclades may have been recruited into the developmental regulatory network underlying ray floret formation.

Previous studies have shown that members of the *CYCLOIDEA2* (*CYC2*) clade are preferentially expressed in ray florets and play central roles in petal elongation and the establishment of dorsoventral floral asymmetry (Chen et al., 2018; Juntheikki-Palovaara et al., 2014; Kim et al., 2008; Li et al., 2026; Shen et al., 2021; Zhang et al., 2024). Tandem duplication of multiple *CYC2* copies has also been reported on chromosome 2 in *Tagetes erecta* (Zhu et al., 2023). We therefore investigated the genomic organization of the *CYC2* locus across 35 Asteraceae species. Surprisingly, we found that the highly expressed *CYC2* genes were all located in collinear genomic blocks, with *CYC2b*, *CYC2c*, *CYC2d*, *CYC2e*, and *CYC2g* consistently forming a tandem gene cluster across all examined species, with a preference of higher expression levels in ray than disk floret, suggesting strong conservation of their genomic organization and potentially conserved biological functions (Table S4-S5 and Figure S3). The size of this cluster varies from 21 kb to 2.3 Mb (Table S4). Because many published Asteraceae genomes have limited contiguity, we examined whether the complete *CYC2* cluster was assembled within a single contig. In most species, the entire cluster was located within a single contig, supporting the conservation of this tandem duplication. However, several *Chrysanthemum* genomes contained fragmented *CYC2* regions across multiple contigs, preventing accurate characterization of this locus. Therefore, a high-quality chromosome-scale genome assembly was required to resolve the structure and evolution of the *CYC2* cluster in this genus.

To this end, we generated a chromosome-scale, haplotype-resolved genome assembly for *Chrysanthemum lavandulifolium* (2n = 2x = 18), whose capitulum is consisted of a single outer whorl of ray florets and multiple inner whorls of disc florets. Therefore, it provides an excellent model for studying capitulum evolution and floral diversification (Figure 1A). Using long-read sequencing and haplotype-resolved assembly approaches, we anchored ∼6.00 Gb of sequences onto 18 pseudochromosomes, representing 98.94% of the 6.06 Gb assembled genome (Table S6 and Figure S4). The longer chromosome from each homologous pair was selected to generate a representative reference genome for downstream analyses. The assembly showed substantial improvements over the previously published scaffold-level genome of *C. lavandulifolium* G1 line (Wen et al., 2022), with contig and scaffold N50 values of 30 Mb and 345 Mb, respectively (Table S7-S8 and Figure S5). Long Terminal Repeat (LTR) retrotransposons were the most abundant repetitive elements, accounting for approximately 72.19% of the genome, with Copia and Gypsy elements representing 35.41% and 27.45%, respectively (Table S9). The genome showed high completeness and accuracy, with a consensus quality (QV) of 73.54, a Benchmarking universal single-copy orthologs (BUSCO) completeness of 97.5%, and an LTR Assembly Index (LAI) score of 25.42 (Table S7 and S10), indicating of a high-quality genome assembly.

**Figure 1.**
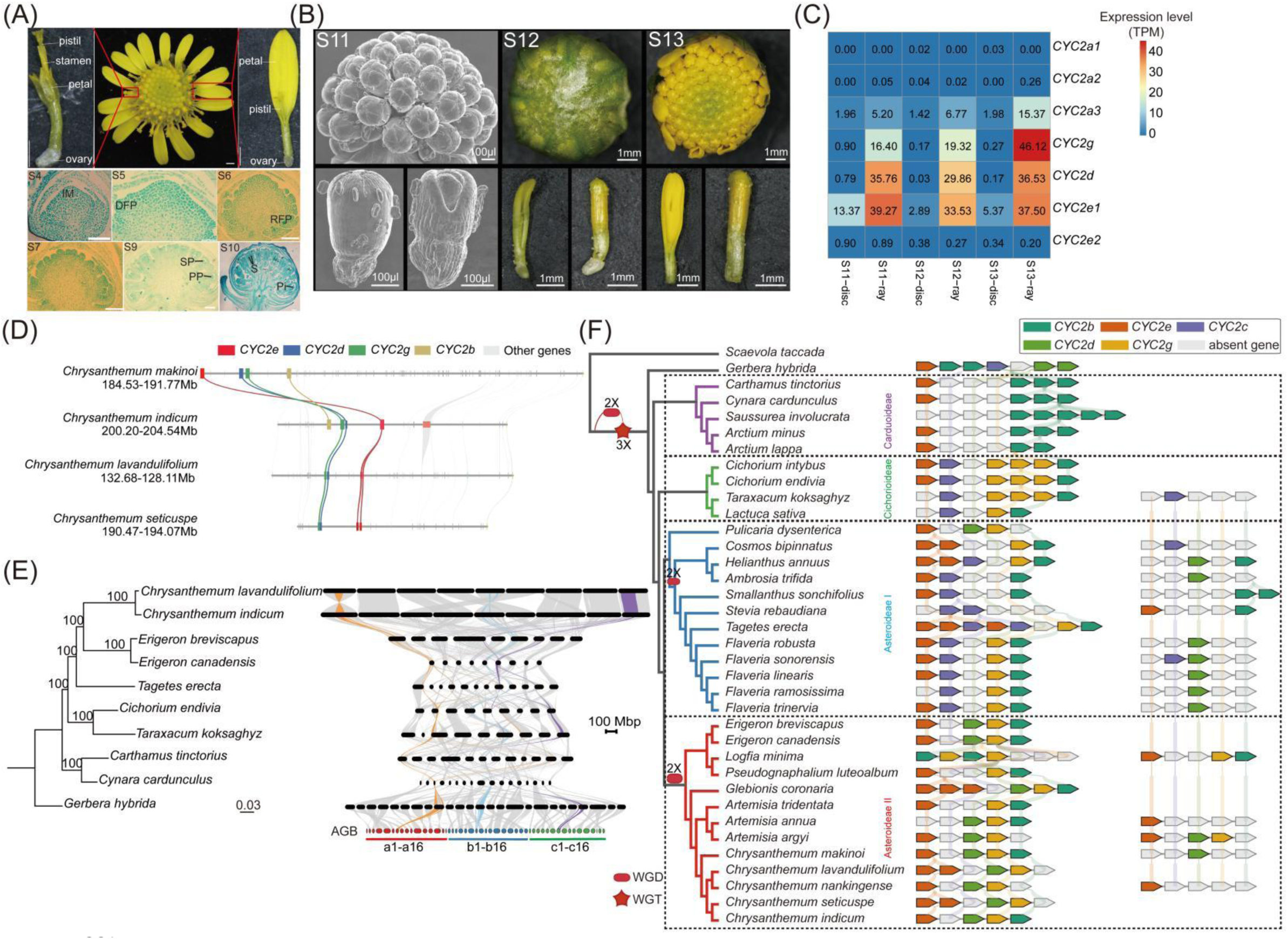
Evolution of a conserved tandem *CYC2* genes cluster in Asteraceae. (A) Morphology and developmental stages of *Chrysanthemum lavandulifolium*. The upper panels show the overall structure of a mature capitulum and the morphology of its two floret types: peripheral ray florets and central disc florets. Ray florets are bilaterally symmetrical and form the showy outer region of the capitulum, whereas disc florets are radially symmetrical and occupy the central reproductive region. The lower panels show paraffin sections of capitula at successive developmental stages. Stages were assigned according to the capitulum morphogenesis framework described by Wen et al. (2019), covering involucre differentiation, floret primordium initiation, corolla primordium differentiation and subsequent floret development. IM: Inflorescence meristem; DFP: Disc florets primordia; RFP: Ray florets primordia; SP: Stamen primordia; S: Stamen; PP: Pistil primordia; Pi: Pistil. Scale bars: 1 mm in the upper panels and 100 μm in the lower panels. (B) Ray florets and disc florets samples used for transcriptome sequencing. Stages S11-S13 were defined in this study as later stages following S10, representing subsequent floret elongation and maturation. (C) Expression profiles of *CYC2* genes in ray and disc florets across developmental stages. Expression levels are shown as transcripts per million (TPM). The profiles reveal stage-dependent and floret-type-biased expression of individual *CYC2* paralogs. (D) Synteny of the *CYC2* gene cluster across four *Chrysanthemum* species. *CYC2e*, *CYC2d*, *CYC2g* and *CYC2b* are highlighted in red, blue, green and yellow, respectively. Grey links connect neighboring collinear genes. (E) Conservation of *CYC2* genes organization across Asteraceae based on the ancestral genome block synteny framework. Left, Phylogenetic relationships of ten representative Asteraceae species spanning major lineages. Bootstrap support values are shown at the nodes. Right, Macrosynteny across the ten Asteraceae genomes shown as a ribbon plot. Tandemly duplicated *CYC2* loci within AGB7-a are highlighted in dark orange, whereas their homologous loci within AGB7-b and AGB7-c are shown in skyblue and purple, respectively. White nodes represent genes located within synteny-inferred genomic intervals. Connecting lines indicate conserved collinearity among species. (F) An evolutionary model summarizing the formation process of the tandem *CYC2* genes cluster in Asteraceae. Whole genome duplication (WGD) events are indicated by red rectangles, and whole genome triplication (WGT) events are indicated by red pentagons. Branch colors indicating subfamily affiliation: purple for Carduoideae, green for Cichorioideae, blue for Asteroideae I, and red for Asteroideae II. The green, orange, purple, yellow, and light green arrows indicate *CYC2b*, *CYC2e*, *CYC2c*, *CYC2g*, and *CYC2d*, respectively. The white arrows represent the corresponding genes that are absent from the genome.

Using this improved genome resource, we identified a complete *CYC2* gene cluster within a 130.96–131.67 Mb interval on chromosome 7 of *C. lavandulifolium*. This haplotype-resolved assembly reconstructed the entire *CYC2* cluster within a single contig, providing a robust framework for investigating its duplication history. To further explore the role of this cluster in capitulum differentiation, we generated transcriptomes from ray and disc florets at multiple developmental stages (Figure 1B and Figure S6 A-C). Comparative expression analysis identified 84 genes consistently upregulated and 252 genes consistently downregulated in ray florets compared with disc florets (Figure S6 D-E). GO enrichment analysis showed that the up-regulated genes were significantly enriched in developmental processes, including regulation of morphogenesis of a branching structure, regulation of secondary shoot formation, and regulation of plant organ formation (Figure S6F). In contrast, the down-regulated genes were enriched in processes related to cellular morphogenesis and pollen development, including cellular component assembly involved in morphogenesis and pollen wall assembly (Figure S6F). Notably, three *CYC2* genes located within the cluster, *CYC2c*, *CYC2d*, and *CYC2e*, showed significantly elevated expression in ray florets, with *CYC2c* and *CYC2d* displaying strong ray floret-specific expression and being barely detectable in disc florets (Figure 1C). These results provide further evidence that the *CYC2* gene cluster plays a central role in regulating ray and disc floret differentiation.

To further trace the evolution of this single tandem duplicated *CYC2* locus in *Chrysanthemum*, we performed a local synteny analysis of the *CYC2* tandem duplication region across four Chrysanthemum genomes, which revealed highly conserved collinearity of both the *CYC2* genes and their flanking genes, suggesting that this tandem cluster has been stably inherited during *Chrysanthemum* evolution (Figure 1D). To investigate the evolutionary origin of the *CYC2* tandem gene cluster, we further selected representative species spanning the major lineages of angiosperms, including the basal angiosperm *Amborella trichopoda*, the basal eudicot *Aquilegia coerulea*, rosids (*Vitis vinifera* and *Glycine max*), asterids outside Asteraceae (*Antirrhinum majus*, *Mimulus verbenaceus*, and *Solanum lycopersicum*), and representative Asteraceae species (*Artemisia annua*, *Helianthus annuus*, and *Chrysanthemum lavandulifolium*) (Table S11). Comparative phylogenomic analysis across these representative angiosperm genomes identified a pronounced, lineage-specific expansion of the *CYC2* subfamily along the Asteraceae stem lineage (Figure S7A and S7B). All three sampled Asteraceae genomes exhibited substantially expanded *CYC2* repertoires (*Helianthus annuus*, n = 9; *Chrysanthemum lavandulifolium*, n = 7; *Artemisia annua*, n = 5), relative to other core eudicots, which consistently retain ≤3 copies (e.g., *Arabidopsis thaliana*, *Vitis vinifera*, *Solanum lycopersicum*, and *Antirrhinum majus*). Although the soybean genome also contains six *CYC2* copies, these genes are dispersed across different chromosomes rather than organized as a tandem array (Figure S7C). In contrast, *CYC2b*, *CYC2c*, *CYC2d*, *CYC2e*, and *CYC2g* consistently form tandem clusters across all examined Asteraceae species. These observations suggest that, unlike the copy number expansion found in soybean, the tandem organization of expanded *CYC2* genes may represent an evolutionary feature specific to the Asteraceae.

To investigate the evolutionary origin of the *CYC2* gene cluster in Asteraceae, we analyzed 10 chromosome-level genome assemblies representing diverse lineages across the family. This dataset includes the recently sequenced genome of *Gerbera hybrida* (Wen et al., 2026), representing an early-diverging lineage within Asteroideae and providing additional phylogenetic context for tracing *CYC2* evolution. Although chromosome-level genome assemblies from the basal Barnadesioideae lineage are currently unavailable, the recently established Asteraceae Genome Blocks (AGBs) framework provides a synteny-based approach to reconstruct the ancestral genomic context of gene evolution across Asteraceae (Feng et al., 2026). The AGB framework defines 48 homologous genomic blocks derived from ancient paleohexaploidization, organized into three sets of corresponding blocks (a, b, and c), which are retained across most extant Asteraceae genomes and provide a comparative framework for tracing genomic evolution. Using the AGB framework, we found that *CYC2* family evolution is tightly associated with ancient genomic architecture (Fig. 1E). The *CYC2b-e,g* genes form a highly conserved tandem gene cluster located within the AGB7-a block, with conserved gene order maintained across diverse Asteraceae lineages despite extensive chromosomal rearrangements (Table S12). In contrast, *CYC2a* is consistently located within the corresponding AGB7-c-derived block. The association of distinct *CYC2* paralog lineages with different AGB7 homologous blocks indicates that major *CYC2* lineages were already partitioned within the ancestral genomic framework established after the paleohexaploidization event. The conserved localization of the *CYC2b-e,g* tandem cluster within the AGB7-a block across extant Asteraceae genomes supports the hypothesis that this cluster originated early in Asteraceae evolution and was subsequently retained throughout diversification, representing an ancient genomic feature associated with the conserved evolution of the AGB7 genomic region.

Within this tandemly duplicated region, we observed that several *CYC2*-like copies have undergone functional loss or complete deletion in specific lineages or capitulum types (Fig. 1F). For instance, members of the *CYC2c*, *CYC2d*, and *CYC2g* clades were entirely absent from Carduoideae, a finding that correlates with the lack of bilaterally symmetrical ray florets in this lineage. Similarly, *Stevia rebaudiana* lacks both *CYC2g* and *CYC2d*; although three *CYC2c* copies were identified, transcriptome analyses revealed they are transcriptionally silent (Fig. S3). The specific and elevated expression of *CYC2c*, *CYC2d*, and *CYC2g* in ray florets, together with the recurrent gains and losses of ray florets across Asteraceae, suggests that these genes represent key components of the regulatory framework underlying the origin and evolutionary diversification of ray floret identity. Intriguingly, we found evidence of functional redundancy within the cluster. In Asteroideae clade II, no *CYC2c* genes were identified (Fig. 1F), while *CYC2d* was absent in the tribe Cichorieae. Despite these specific losses, species in both groups retain the ability to form normal ray florets, suggesting that *CYC2c*, *CYC2d*, and *CYC2g* may functionally compensate for one another during floral development.

To elucidate the genomic mechanisms underlying gene loss within the tandem *CYC2* region, we evaluated whether transposable elements (TEs) are preferentially associated with *CYC2* genes. Because TE insertions can disrupt gene structure, promote unequal recombination, or contribute to local genomic instability, we quantified TE overlap within *CYC2b*-*e*, *g* gene regions and compared these values with those of randomly selected genomic genes from the same chromosomal background. A total of 172 *CYC2b-e, g* genes were included in the analysis. As a genomic background control, 172 non-*CYC2* genes were randomly sampled from the same chromosomes. Pairwise one-way ANOVA comparisons between each *CYC2* subgroup and the random gene set revealed no significant differences in TE density between *CYC2b-e, g* genes and the randomly selected genes (Fig. S8A). We further examined the composition of TEs overlapping *CYC2* and random gene regions. In random genes, LTR retrotransposons were the dominant TE class, representing 67.2% of the total TE-overlap length, whereas DNA transposons accounted for 25.9%. In contrast, most *CYC2* subgroups were characterized by a predominance of DNA transposons. DNA transposons accounted for 54.2% of TE overlaps in *CYC2b*, 68.1% in *CYC2d*, 60.5% in *CYC2e* and 71.6% in *CYC2g* genes, whereas *CYC2c* genes showed a distinct pattern dominated by LTR retrotransposons (Fig. S8B). This shift from LTR-dominated TE composition in random genes to DNA-transposon-dominated composition in most *CYC2* genes suggests that *CYC2* loci may have experienced a different spectrum of TE insertions from the genomic background. Given that DNA transposons can directly interrupt gene sequences and promote local rearrangements, their enrichment within *CYC2* gene regions provides a potential mechanism for gene degeneration and loss within the tandem *CYC2* region.

In summary, we identified a conserved tandem *CYC2* gene cluster that is broadly distributed across Asteraceae. Phylogenetic analyses indicate that a single-copy ancestral *CYC2* gene, as seen in the outgroup Campanulaceae, underwent two rounds of whole-genome triplication, giving rise to the primary *CYC2* paralogs in Asteraceae. This gene cluster was established early in the evolutionary history of the family. Subsequent insertions of DNA transposable elements (TEs) may have contributed to the loss of function and neofunctionalization of some *CYC2* copies. These changes may have facilitated the diversification of capitulum architecture and floral morphology in Asteraceae.

## Acknowledgments

We apologize to colleagues whose work could not be fully cited owing to space constraints. This work was financially supported by grants from the National Natural Science Foundation of China to B.D. (32673535; 32122078). We are grateful to Dr. Haiyan Yuan from the Institute of Botany, Jiangsu Province and Chinese Academy of Sciences for providing the *Stevia rebaudiana* seedlings. We also thank Dr. Jiayu Xue and Ms. Xuefen Wei of Nanjing Agricultural University for their assistance with the early version of the genome assembly. We would like to acknowledge Dr. Eric Schranz and Dr. Tao Feng of Wageningen University and Research for assisting with synteny analysis across Asteraceae species. We are grateful to Dr. Yuehua Ma (Central Laboratory of the College of Horticulture, Nanjing Agricultural University) for assistance in using the Scanning Electron Microscope FlexSEM 1000II (Hitachi). This project was supported by the Bioinformatics Center of Nanjing Agricultural University.

## Author Contributions

D.Z. and B.D. planned and designed the research; D.Z., Y.G., Y.Q., Y.S., J.Z., Y.L., Y.H. performed most of the data collection and bioinformatics analyses; F.C. and S.C. provided insightful advice, discussions and support throughout. And Z.D. and B.D. wrote the manuscript with input from all authors.

## Data and code availability

All raw data are available in the National Genomics Data Center (https://ngdc.cncb.ac.cn/) under project number PRJCA033117. The assemblies and annotations of *Chrysanthemum lavandulifolium* have been deposited in the Chrysanthemum Genome Database (CGD) (https://cgd.njau.edu.cn/asteraceae/download/downloadPage). The analysis code is available at https://github.com/zhangdeng1992/genome-of-Chrysanthemum-lavandulifolium.

## Materials and methods

### Plant materials

The ***C. lavandulifolium* genomic DNA sample was collected** at the *Chrysanthemum* Germplasm Resource Preserving Centre (Nanjing Agricultural University, Nanjing, Jiangsu, China) for genomic sequencing. All of the samples were collected from a single individual *C. lavandulifolium* plant. Genomic DNA was extracted from young leaves using a standard CTAB protocol for Illumina short-read sequencing, PacBio long-read sequencing, and Hi-C library construction and sequencing. Fresh samples were collected, immediately frozen in liquid nitrogen, and then stored at -80°C for subsequent nucleic acid extraction. Replicates were obtained from separate clonal plants.

### Genome sequencing and library construction

For PacBio long-read sequencing, three approximately 20-kb SMRTbell libraries were prepared using the SMRTbell Express Template Preparation Kit 1.0 following the PacBio 20-kb protocol (https://www.pacb.com/). Sequencing was performed on the Sequel II platform in the circular consensus sequencing (CCS) mode. A total of ∼90 Gb of PacBio HiFi read data was generated using CCS (v4.0, https://github.com/PacificBiosciences/ccs). Two Hi-C libraries were constructed by chromatin extraction, digestion with MboI **enzyme**, purification, and fragmentation. A total of ∼150 Gb of raw reads was obtained on the MGISEQ-2000 platform in paired-end mode.

### Genome assembly and assessment

We used a haplotype-resolved assembly strategy to integrate the genomic HiFi reads with Hi-C data to produce haplotype-resolved contigs using hifiasm v0.25.0 (Cheng et al., 2021) with the parameters ‘-u1 -l1 -s 0.75’. Before chromosome assembly, the Hi-C data were mapped to the raw alternate assemblies from hifiasm using BWA v0.7.17 (Li, 2013). The uniquely mapped data were subjected to YaHS v1.2.2 (Zhou et al., 2023) scaffolding with the default parameters and then partitioned into 9 groups, representing 9 pseudo-chromosomes.

### Repeat identification and gene annotations

After generating chromosome-level assemblies for the two haplotypes, both genomes were subjected to genome annotation. EDTA v2.2.2 (Ou et al., 2019) was used to **de novo identify TEs in the two haplotypes genomes**. Inpactor2 v1.0 (Orozco-Arias et al., 2023) was used to recover full-length LTR retrotransposons in the two genomes and to classify them at the lineage level. EDTA and Inpactor2 libraries were merged and clustered using CD-HIT v4.8.1 (Fu et al., 2012). Repetitive elements were identified using the merged and clustered libraries and masked with RepeatMasker v4.1.7 (Smit et al., 2015) using the softmasking option.

Gene structural annotation was performed by BRAKER pipeline v3.0.6 (Brůna et al., 2021), which combines evidence from de novo prediction, transcriptome data, and protein homology. Following the initial gene prediction by BRAKER, the resulting output was further processed using EviAnn v2.0.3 (Zimin et al., 2026) for improved evidence-based annotation. Annotation completeness was assessed with BUSCO v5.4.7 (Simão et al., 2015).

### Comparative genomics analysis

Hierarchical Orthologous Groups (HOGs, gene families) were inferred using FastOMA v1.2.0 (Majidian et al., 2025), with a species tree reconstructed by OrthoFinder v2.5.5 (Emms and Kelly, 2019) as the guide phylogeny, with the parameters ‘-M msa -S diamond’. We selected single-copy HOGs across all species as the seed sequences. The protein sequences of genes in each HOG were aligned using MAFFT v7.475 (Katoh and Standley, 2013). The alignments were then further trimmed with Gblocks v0.91b (Castresana, 2000) using the parameters ‘-t=p -b4=2 -b5=h’. Using the concatenated alignment of these groups, we inferred a phylogenetic tree using RAxML-NG v2.0.0 (Kozlov et al., 2019) with the LG+G8+F model and with 200 bootstrap replicates. Finally, the phylogenetic trees were visualized with iTOL v6 (Letunic and Bork, 2024). To identify genes present in Asteroideae but absent from Carduoideae, we reconstructed the genomes of the most recent common ancestors (MRCAs) of Asteroideae and Carduoideae using the get_ancestral_genome_by_name() function in pyHam (Train et al., 2019). We then compared the HOGs inferred for the Asteroideae and Carduoideae MRCAs and identified HOGs present in the Asteroideae MRCA but absent from the Carduoideae MRCA. Candidate HOGs were further identified using HMMER v3.3.2 (Eddy, 2011) searches against the corresponding HMM profiles with an E-value cutoff of 1e-5.

### Phylogenetic analyses of the *CYC2* genes

*CYC2* homologs were identified based on hierarchical orthogroups (HOGs), and the corresponding protein sequences were used to construct the *CYC2* phylogenetic tree. We aligned these sequences using MAFFT v7.475 (Katoh and Standley, 2013) with the ‘--localpair’ parameter, and corresponding coding sequence alignments were trimmed using trimAl v1.5 (Capella-Gutiérrez et al., 2009). Gene family trees were reconstructed using RAxML-NG v2.0.0 (Kozlov et al., 2019) with the LG+G8+F model and with 200 bootstrap replicates. Abnormally long branches that significantly inflated tree diameter were detected using TreeShrink v1.3.9 (Mai and Mirarab, 2018), with the false positive rate set to 0.1 (option -q). The corresponding sequences were then removed from the alignments, and the phylogenetic analysis was repeated. Finally, the phylogenetic trees were visualized with iTOL v6 (Letunic and Bork, 2024).

### Transcriptome analysis

Available transcriptome datasets of Asteraceae species were downloaded from the NCBI Sequence Read Archive (SRA) (Supplemental Table 6). The clean reads were mapped to the genome of the corresponding species using STAR v2.7.5b (Dobin et al., 2013) with the parameters --twopassMode Basic, --outMultimapperOrder Random, and --outSJfilterReads Unique. The gene expression analyses were performed using the ‘rsem-calculate-expression’ program from RSEM v1.3.3 (Li and Dewey, 2011) after generating indexes for the reference genome and GTF files using ‘rsem-prepare-reference’. Genes with counts > 4 in at least four samples were retained for downstream differential expression analysis using DESeq2 (Love et al., 2014). Differentially expressed genes were selected using the criteria of an absolute log◻fold change ≥ 1 and an adjusted p-value < 0.05. The heatmap of expression levels was plotted using the pheatmap v1.0.12 package (Kolde, 2019).

### Multigenome macrosynteny and microsynteny

Global synteny across the 12 genomes was inferred using a custom script developed by Feng et al. (2026) for Asteraceae synteny-phylogenomic analysis in Asteraceae. The macrosynteny among the four species of the genus *Chrysanthemum* was analyzed using the compara.catalog and graphics.synteny modules of JCVI v1.2.7 (Tang et al., 2024).

### GO annotation and enrichment analysis

Protein sequences were searched against the protein sequences of Araport11 (Cheng et al., 2017) with DIAMOND BLASTP v2.1.24 (Buchfink et al., 2021) in sensitive mode, using an E-value cutoff of 1e-5 and --max-target-seqs 1. For each query sequence, only the best-scoring hit was retained, and GO terms associated with the corresponding Arabidopsis locus were transferred from the TAIR Gene Association File (GAF). GO enrichment analysis was performed using the GOATOOLS v1.6.5 package (Klopfenstein et al., 2018). GO terms were tested using Fisher’s exact test, and P values were adjusted using the Benjamini-Hochberg false-discovery-rate (FDR) procedure.

---

**Supplemental Figure 1.**
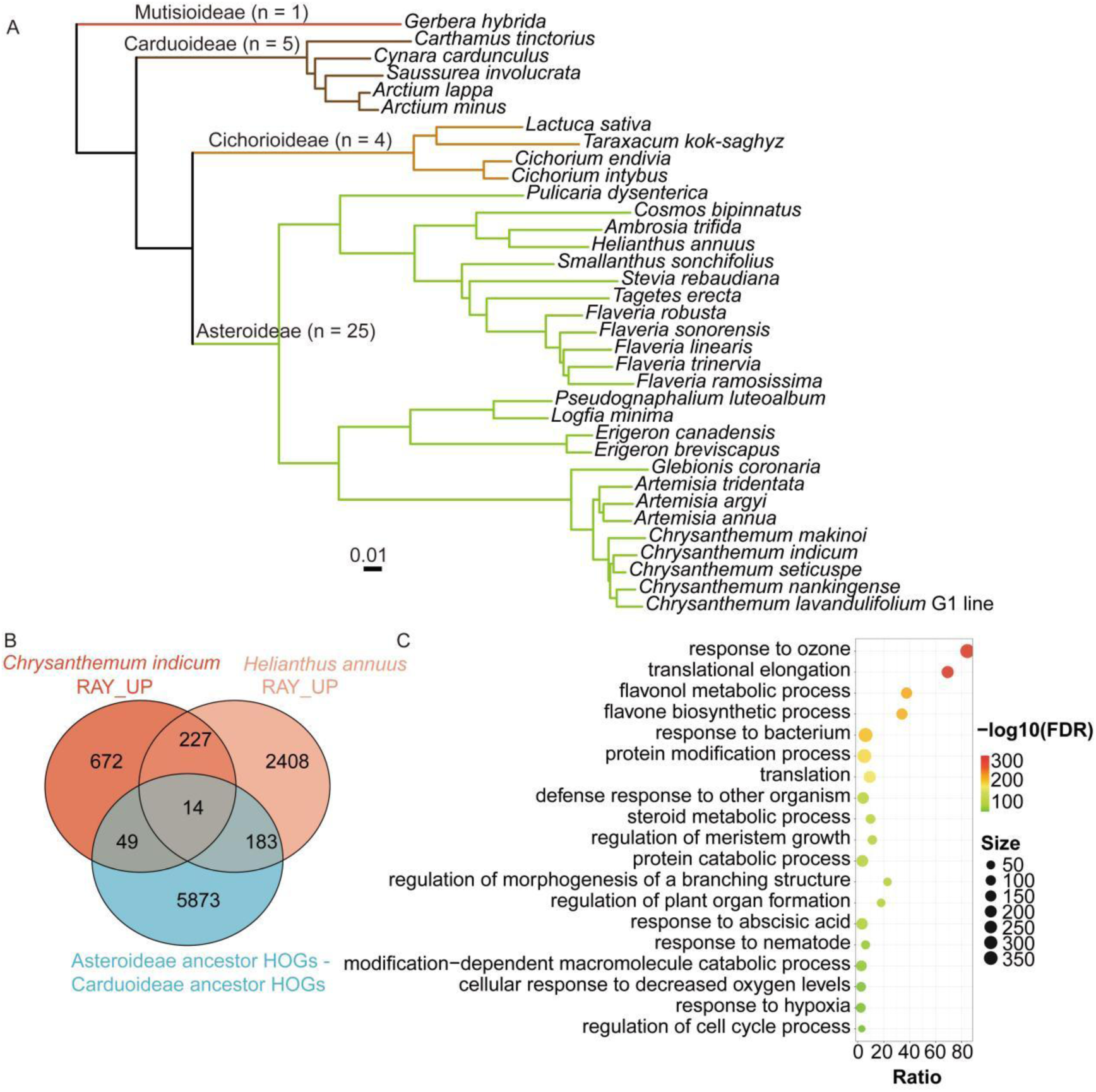
Evolutionary identification of Asteroideae-specific orthologous gene groups associated with ray floret identity. **A.** Maximum-likelihood phylogeny of 35 Asteraceae species. Branch colors indicate subfamily affiliation: Mutisioideae (red), Carduoideae (brown), Cichorioideae (orange), and Asteroideae (green) **B.** Overlap between Asteroideae lineage-specific HOGs and genes upregulated in ray florets of *Chrysanthemum indicum* and *Helianthus annuus*. **C.** GO enrichment analysis of the 14 overlapping HOGs. Bubble size indicates the number of genes associated with each GO term, and color represents enrichment significance as −log10(FDR).

**Supplemental Figure 2.**
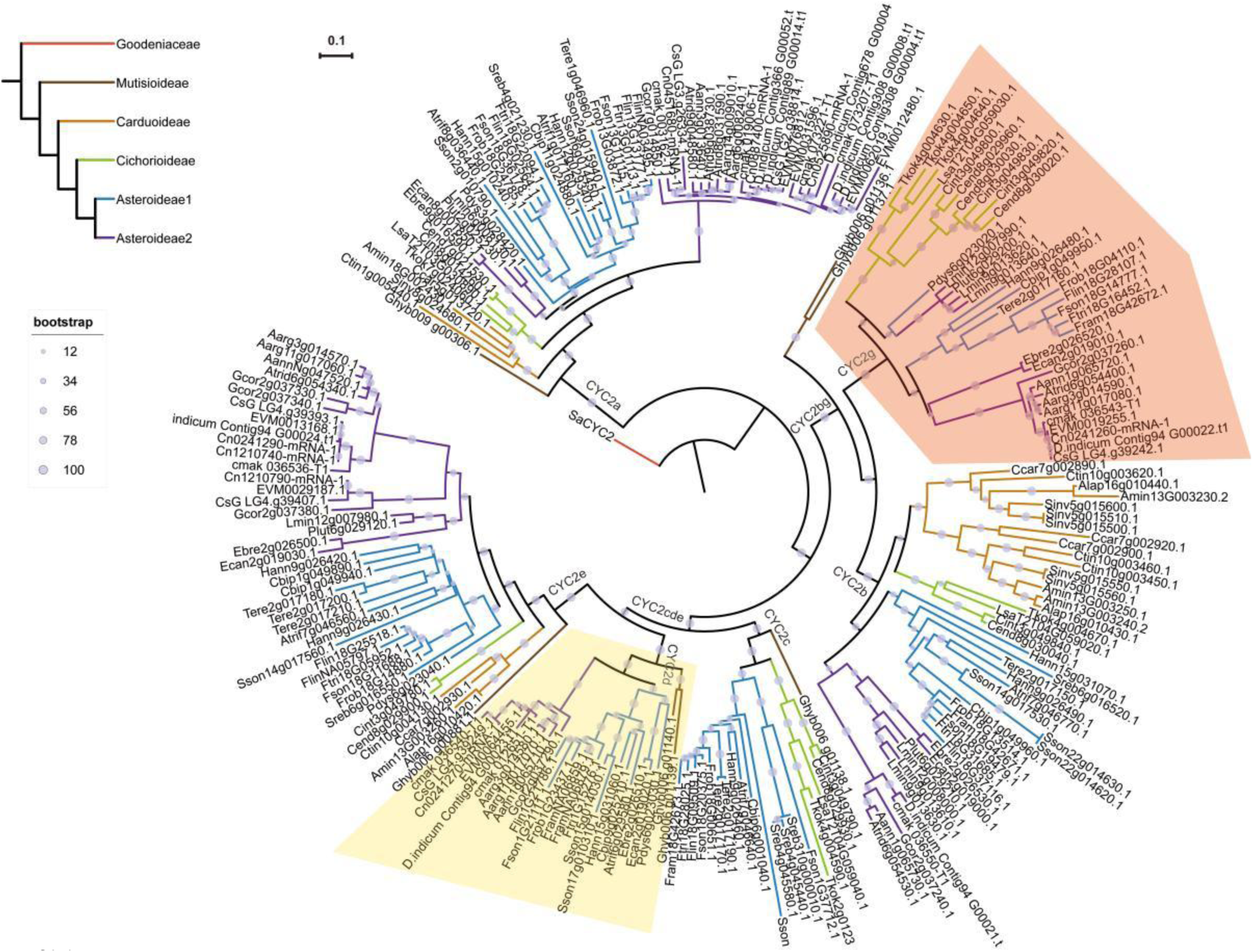
Phylogenetic relationships of *CYC2*-like genes in the Asterales inferred from 230 deduced amino acid sequences. The maximum-likelihood (ML) tree was reconstructed under the LG+G8+F substitution model with 200 bootstrap replicates and rooted using Goodeniaceae sequences as the outgroup. ML bootstrap support values are represented by the size of the circles at internal nodes. The *CYC2d* and *CYC2g* clades are highlighted in red and yellow, respectively.

**Supplemental Figure 3.**
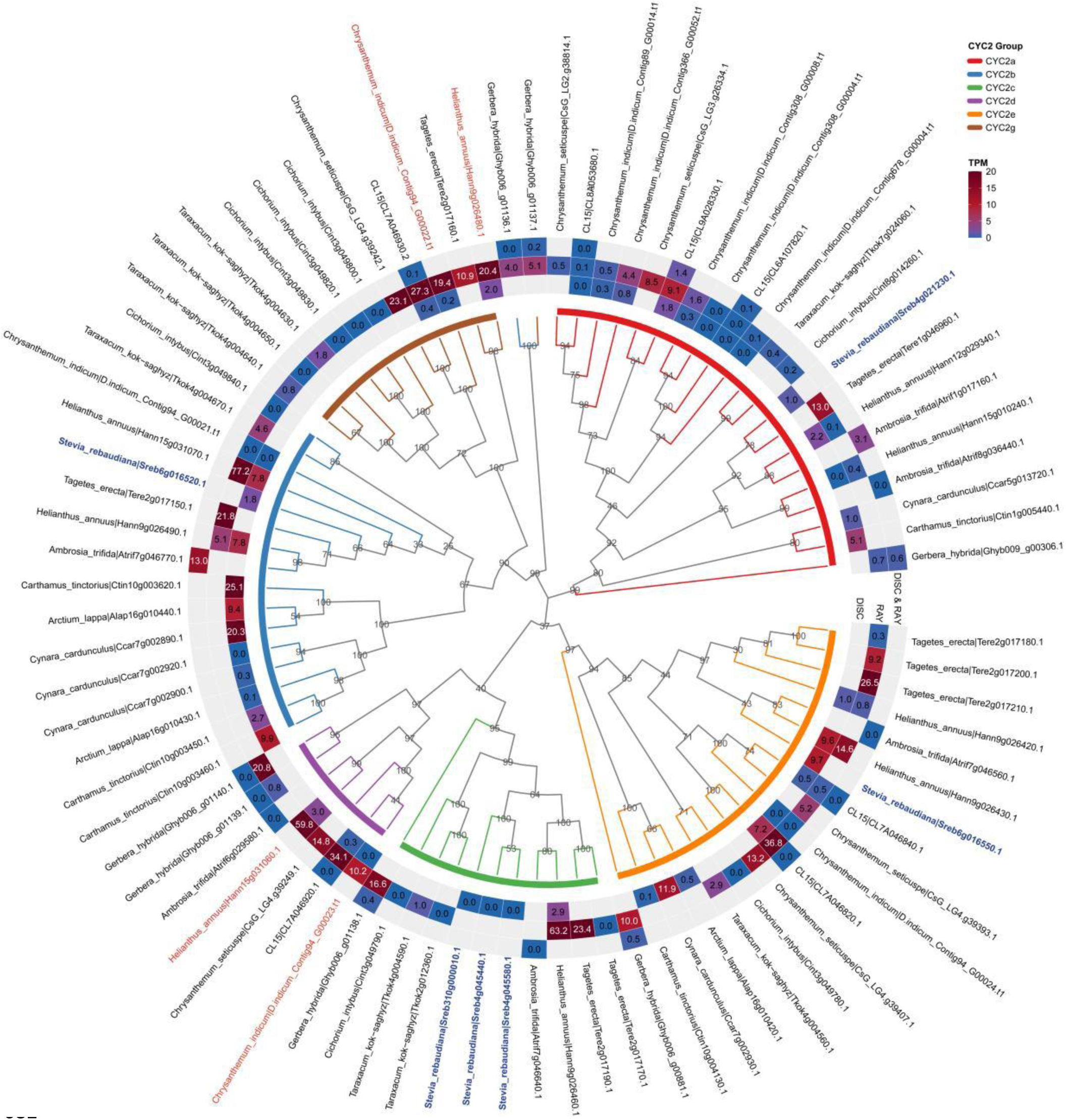
Phylogenetic relationships and expression patterns of *CYC2*-like genes in Asteraceae. A maximum-likelihood phylogenetic tree of *CYC2*-like genes from multiple Asteraceae species is shown in the center. Branches are colored according to major *CYC2* subclades (*CYC2a*-*CYC2e*, *CYC2g*). Bootstrap support values are indicated at internal nodes. Gene labels are formatted as species name | gene ID, where the species name and gene identifier are separated by a vertical bar (“|”). The outer circular tracks display gene expression levels (TPM) across different tissues or floral organs. Heatmap colors represent normalized expression values, with higher expression shown in red and lower expression shown in blue. The *CYC2d* and *CYC2g* genes from *Chrysanthemum indicum* and *Helianthus annuus* are highlighted in red, whereas the *CYC2*-like genes from *Stevia rebaudiana* are highlighted in blue and shown in bold on the phylogenetic tree.

**Supplemental Figure 4.**
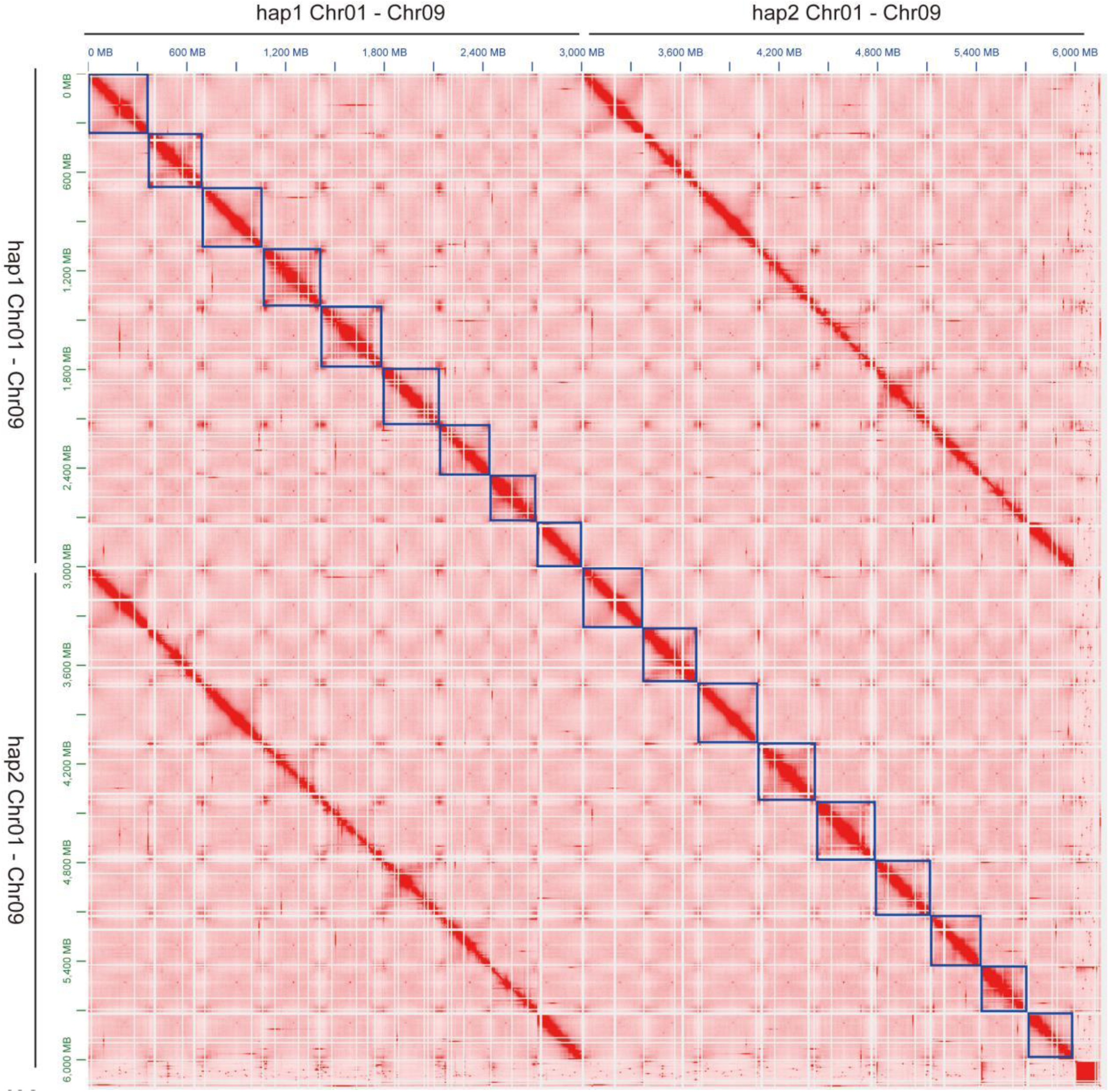
Hi-C contact heatmap of nine homologous chromosome pairs in *Chrysanthemum lavandulifolium*. The heatmap displays genome-wide chromatin interaction frequencies for nine groups of homologous chromosomes (Chr01–Chr09), visualized as a pairwise contact matrix. Interaction intensity is color-coded from white (low contact frequency) to red (high contact frequency). Each block represents intra-chromosomal interactions within and inter-chromosomal interactions between the nine homologous chromosome groups. The diagonal reflects the strength of local chromatin interactions along each chromosome.

**Supplemental Figure 5.**
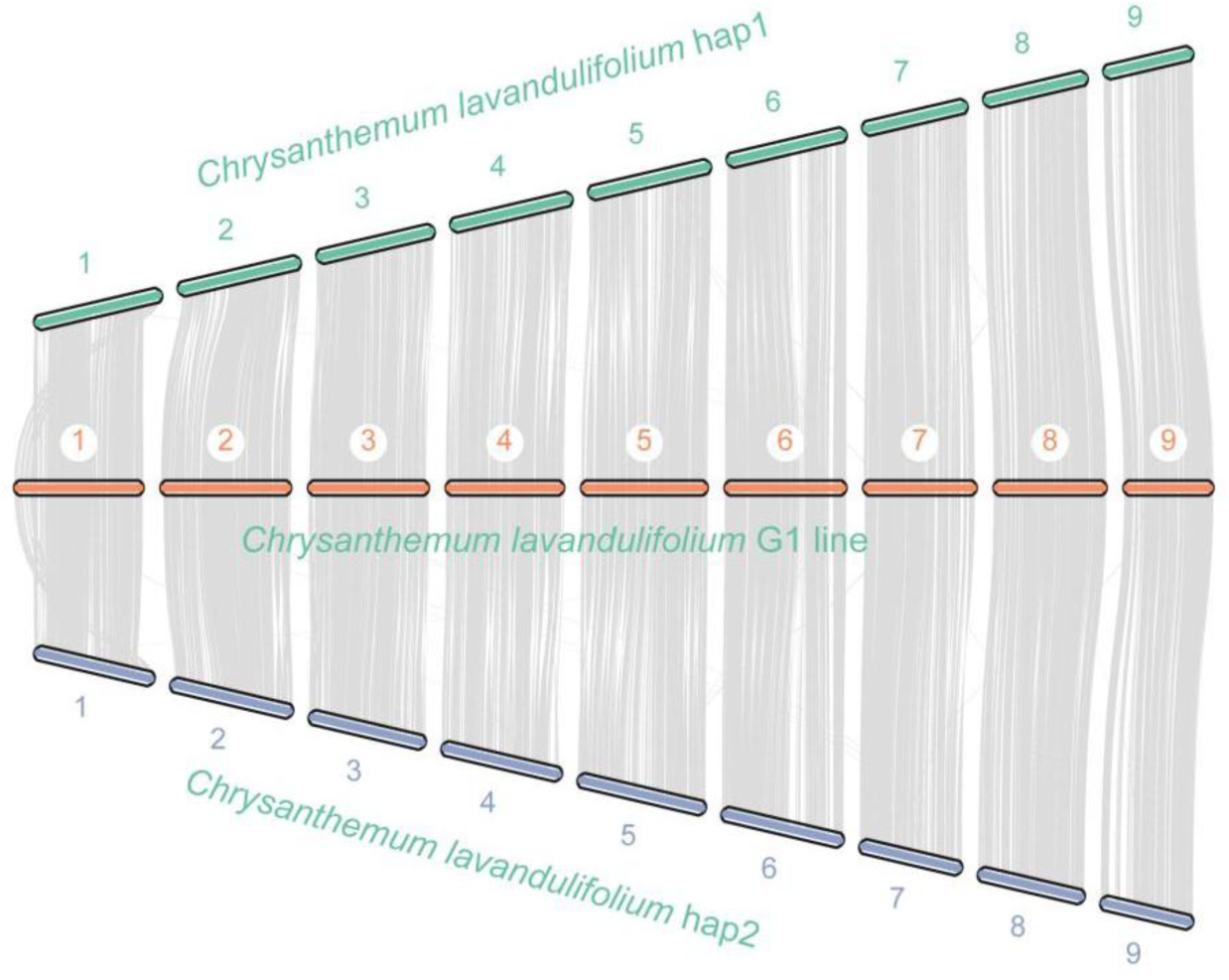
Chromosome-scale synteny between the G1 assembly and two phased haplotypes of *Chrysanthemum lavandulifolium*. Syntenic relationships among the G1 assembly (middle track) and two phased haplotypes (hap1, upper track; hap2, lower track) of *Chrysanthemum lavandulifolium* are shown. Chromosomes are numbered from 1 to 9 and displayed in the corresponding order. Grey ribbons connect collinear gene pairs identified using the JCVI. Extensive chromosome-scale collinearity is observed among the three assemblies, indicating high structural conservation and assembly continuity. Minor local deviations in collinearity likely reflect haplotype-specific structural variation.

**Supplemental Figure 6.**
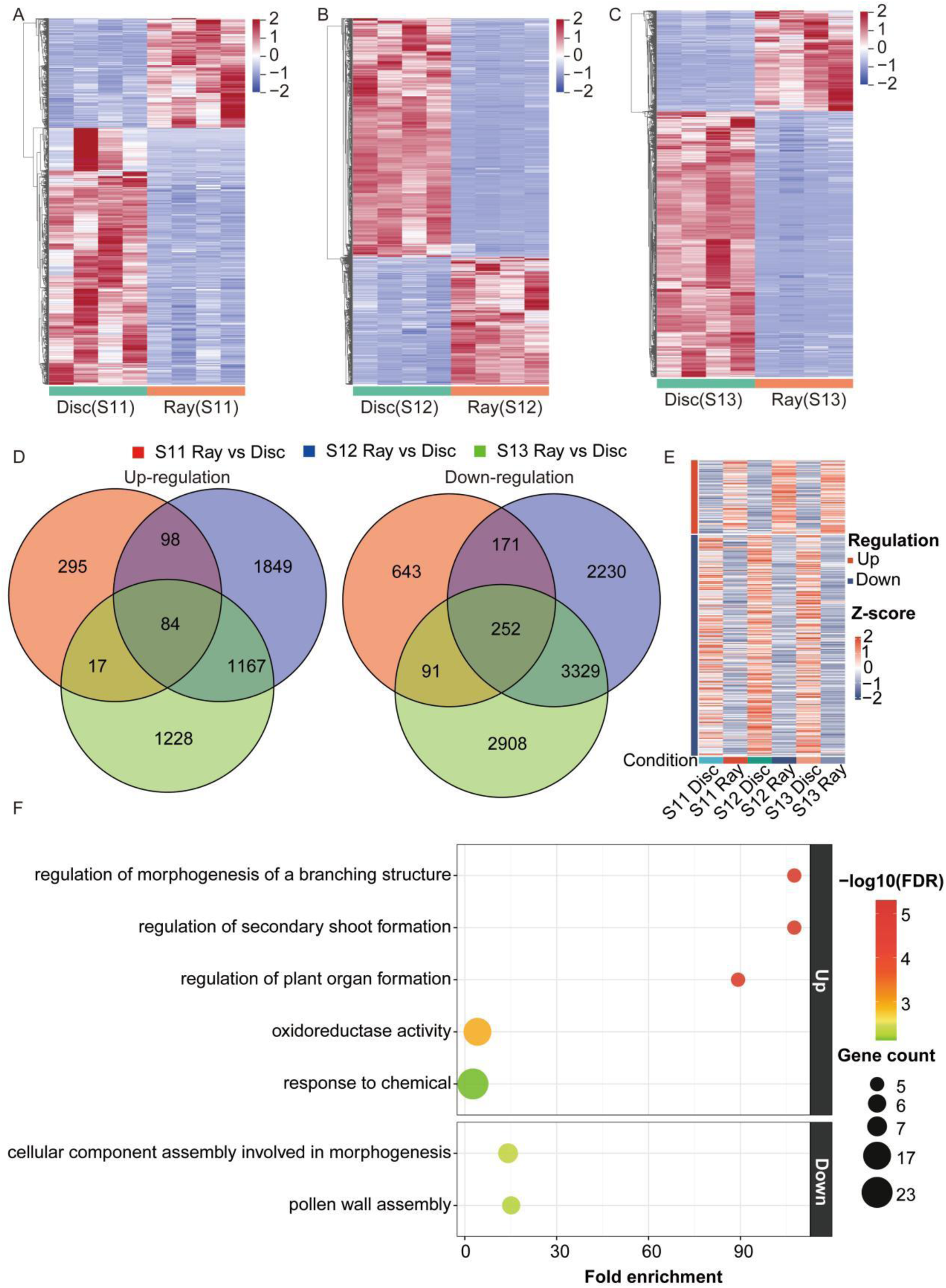
Comprehensive transcriptomic analysis of ray and disc florets of *Chrysanthemum lavandulifolium* across developmental stages. **A**–**C.** Heatmaps showing the differentially expressed genes between ray and disc florets across developmental stages S11 (A), S12 (B), and S13 (C). **D.** Venn diagrams showing the overlap of differentially expressed genes in ray and disc florets across developmental stages S11, S12, and S13. **E.** Heatmap of overlapping differentially expressed genes in ray and disc florets across developmental stages S11, S12, and S13. Expression values were standardized by row using Z-scores. Red and blue indicate relatively high and low expression, respectively, and the left-side annotation indicates up- or down-regulation. **F.** GO enrichment analysis of overlapping differentially expressed genes in ray and disc florets across developmental stages S11, S12, and S13. Bubble size indicates the number of genes assigned to each Gene Ontology biological process term, and color represents significance as −log10(FDR). Only terms at ontology depth 5 with FDR < 0.01 are shown.

**Supplemental Figure 7.**
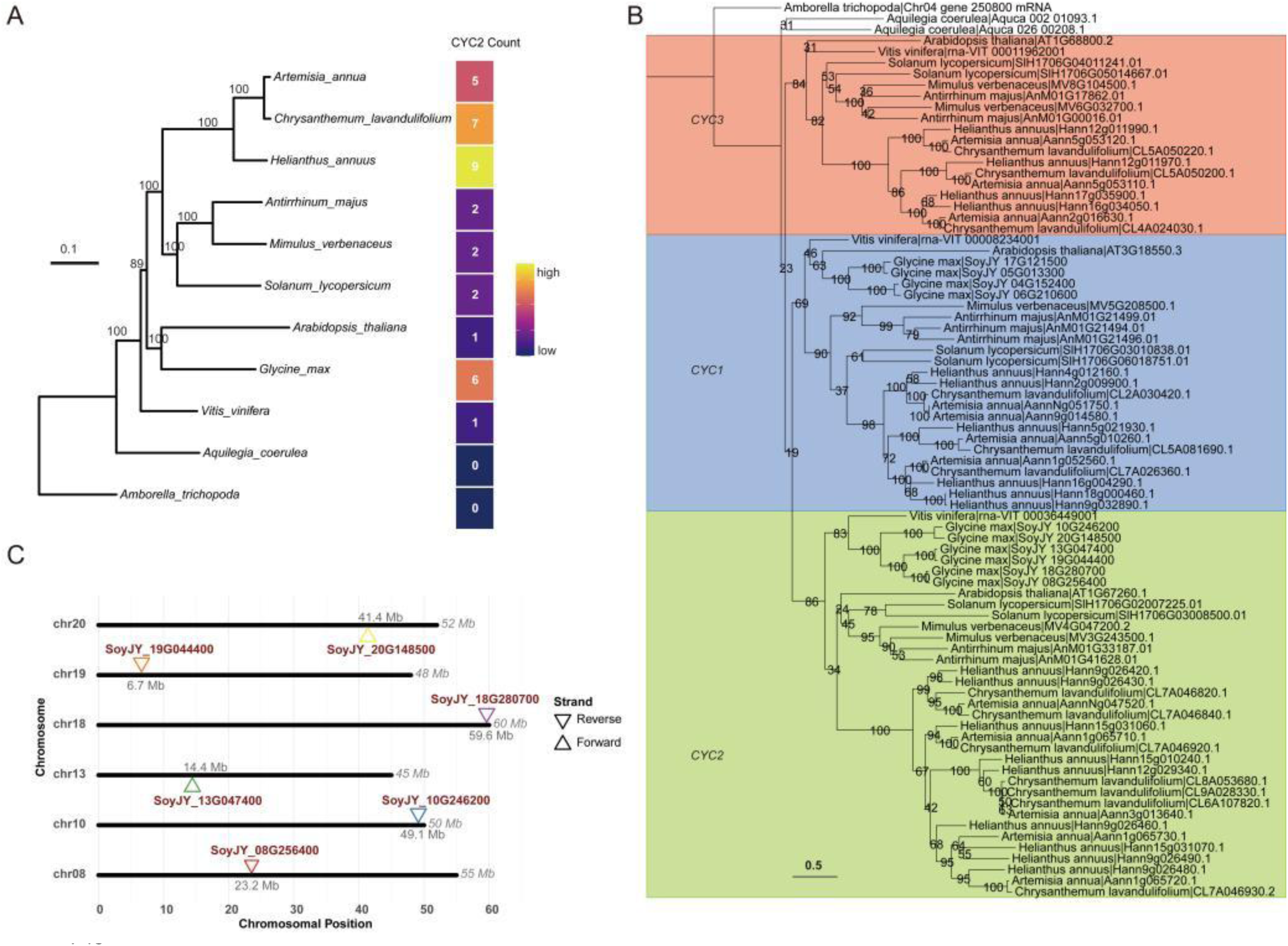
Evolutionary diversification of the *CYC/TB1* gene family and expansion of *CYC2* paralogs in Asteraceae. **A.** Species phylogeny of 11 representative angiosperm species used for comparative analyses. Numbers at nodes indicate bootstrap support values. The adjacent heatmap shows *CYC2* gene copy numbers identified in each species. **B.** Maximum-likelihood phylogeny of *CYC/TB1* TCP family proteins from the 11 representative angiosperm species. Three major clades corresponding to *CYC1*, *CYC2*, and *CYC3* are highlighted by colored backgrounds. Bootstrap support values are indicated at nodes. Protein sequences were aligned using MAFFT v7 with the L-INS-i strategy (--localpair --maxiterate 1000), followed by alignment trimming using trimAl (-gt 0.15). The maximum-likelihood tree was inferred using RAxML-NG under the LG+G8+F substitution model with 200 bootstrap replicates. **C.** Chromosomal locations of *CYC2* homologs in *Glycine max*. Triangles indicate gene positions and orientations on chromosomes. Upward and downward triangles represent forward and reverse strand orientations, respectively.

**Supplemental Figure 8.**
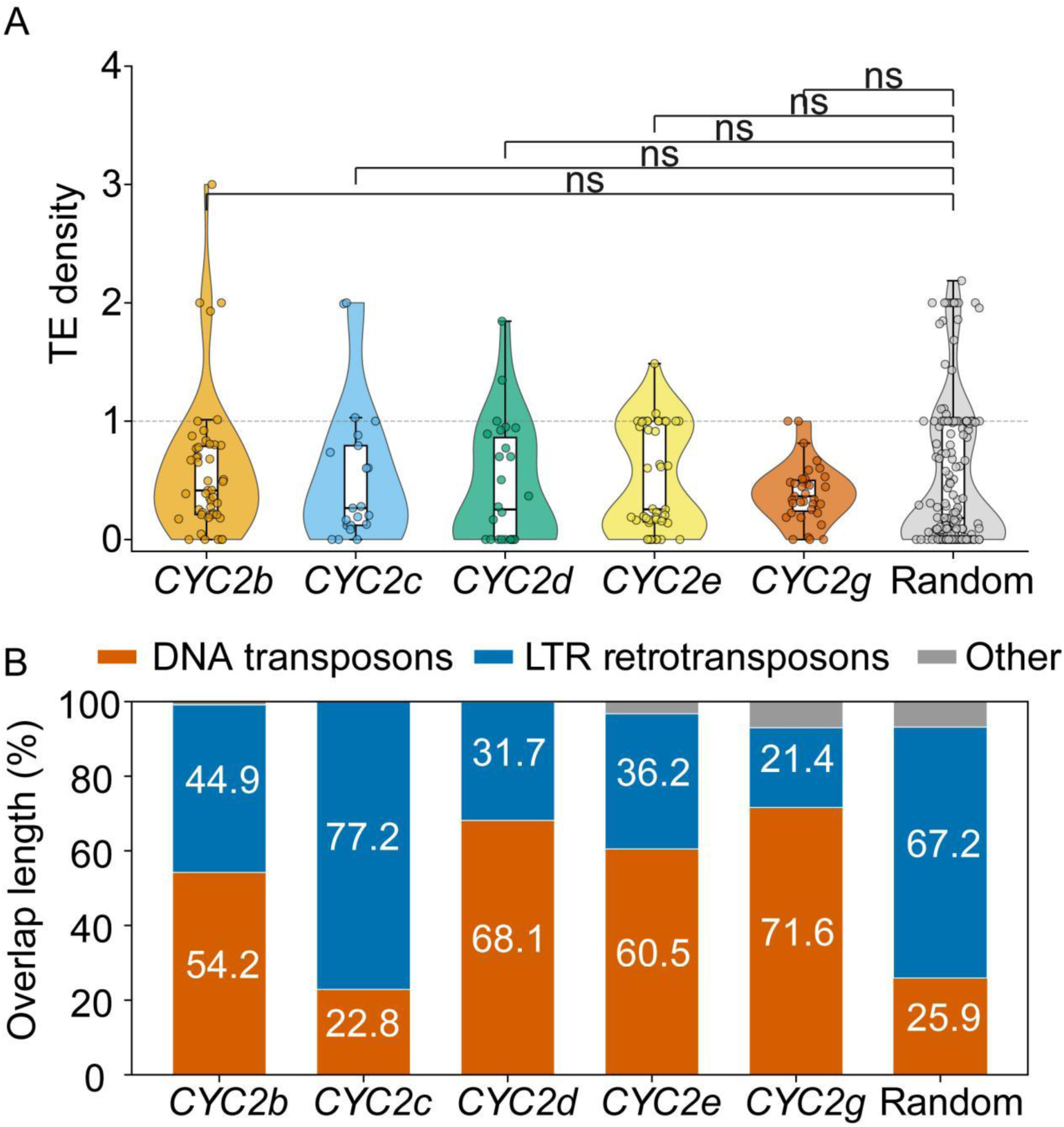
Statistical analysis of transposable elements (TEs) within *CYC2b-e,g* genes. **A.** TE density distribution in *CYC2b-e,g* genes and randomly sampled protein-coding genes. A total of 172 *CYC2b-e,g* genes were included, and 172 non-*CYC2* genes were randomly sampled from protein-coding genes on the same chromosomes as the *CYC2b-e,g* genes. TE density was calculated as the total length of TE overlaps divided by gene length. Violin plots show the distribution of TE density; internal boxplots indicate the median and interquartile range, and individual points represent individual genes. The dashed horizontal line indicates a TE density of 1. Statistical comparisons were performed using one-way ANOVA. Significance labels indicate *P < 0.05 and *ns*, not significant. B. TE class composition of gene-overlapping regions. Bars show the percentage of total TE-overlap length assigned to three TE classes: DNA transposons, LTR retrotransposons, and “Other”. The “Other” category includes MITE, “Unknown”, and non-DNA/LTR TE annotations.

